# Natural selection and genetic diversity in the butterfly *Heliconius melpomene*

**DOI:** 10.1101/042796

**Authors:** SH Martin, M Möst, WJ Palmer, C Salazar, WO McMillan, FM Jiggins, CD Jiggins

**Affiliations:** Department of Zoology, University of Cambridge, Downing Street, Cambridge, CB2 3EJ, UK; Department of Genetics, University of Cambridge, Downing Street, Cambridge, CB2 3EH, UK; Biology Program, Faculty of Natural Sciences and Mathematics, Universidad del Rosario, Carrera 24 No 63C-69, Bogota 111221, Colombia; Smithsonian Tropical Research Institution, Apartado 0843 – 03092, Balboa, Ancón, Panama

**Keywords:** background selection, genetic hitchhiking, recombination rate, selective sweeps, effective population size

## Abstract

A combination of selective and neutral evolutionary forces shape patterns of genetic diversity in nature. Among the insects, most previous analyses of the roles of drift and selection in shaping variation across the genome have focused on the genus *Drosophila*. A more complete understanding of these forces will come from analysing other taxa that differ in population demography and other aspects of biology. We have analysed diversity and signatures of selection in the neotropical *Heliconius* butterflies using resequenced genomes from 58 wild-caught individuals of *H. melpomene*, and another 21 resequenced genomes representing 11 related species. By comparing intra-specific diversity and inter-specific divergence, we estimate that 31% of amino acid substitutions between *Heliconius* species are adaptive. Diversity at putatively neutral sites is negatively correlated with the local density of coding sites as well as non-synonymous substitutions, and positively correlated with recombination rate, indicating widespread linked selection. This process also manifests in significantly reduced diversity on longer chromosomes, consistent with lower recombination rates. Although hitchhiking around beneficial non-synonymous mutations has significantly shaped genetic variation in *H. melpomene*, evidence for strong selective sweeps is limited overall. We did however identify two regions where distinct haplotypes have swept in different populations, leading to increased population differentiation. On the whole, our study suggests that positive selection is less pervasive in these butterflies as compared to fruit flies; a fact that curiously results in very similar levels of neutral diversity in these very different insects.

## INTRODUCTION

Genetic variation within and between populations is shaped by numerous factors. In particular, genetic drift is stronger in smaller populations, such that organisms with larger population sizes should be more diverse under neutral evolution. However, it has long been known that the amount of genetic variation does not always scale as expected with population size, with a deficit of genetic variability in larger populations as compared to the neutral expectation (Lewontin 1974). This has become known as ‘Lewontin’s Paradox’. It is likely that this paradox can be explained by considering the influence of natural selection (Ohta and Gillespie 1996; Leffler *et al.* 2012; Cutter and Payseur 2013; Corbett-Detig *et al.* 2015). Since drift can act to retard selection, natural selection tends to be more efficient in organisms with larger population sizes. Consistent with this, estimated rates of adaptive evolution are often greater for smaller organisms with larger population sizes. For example, it has been estimated that over 50% of amino acid substitutions between fruit fly species are driven by positive selection (Sella *et al.* 2009; Messer and Petrov 2013), but in humans less than 15% of recent amino acid substitutions appear to have been driven by selection (Eyre-Walker 2006; Messer and Petrov 2013). Considering the relative importance of natural selection and genetic drift in maintaining genetic diversity has important implications for explaining current patterns of biodiversity and predicting future adaptive potential (Gillespie 2001; Leffler *et al.* 2012).

The solution to Lewontin’s Paradox appears to lie in the influence of natural selection on linked sites. Selection acting on one locus can cause the removal of genetic variation at physically linked, neutral loci. This can occur either through fixation of beneficial alleles (‘hitchhiking’) (Maynard Smith and Haigh 1974), or by purging of deleterious alleles (‘background selection’) (Charlesworth *et al.* 1993). Both of these processes have more pronounced effects in genomic regions of lower recombination rate. The importance of linked selection is supported by a positive correlation between recombination rate and neutral genetic diversity, first and most thoroughly studied in *Drosophila melanogaster* (Begun and Aquadro 1992; Mackay *et al.* 2012; McGaugh *et al.* 2012; Langley *et al.* 2012; Campos *et al.* 2014), and subsequently observed in other taxa, including humans (Nachman *et al.* 1998; Payseur and Nachman 2002; McVicker *et al.* 2009; Lohmueller *et al.* 2011), yeast (Cutter and Moses 2011), mice, rabbits (Nachman and Payseur 2012), and chickens (Mugal *et al.* 2013). A major recent advance has come from a population genomic analysis of 40 species, which showed not only that this phenomenon is widespread in plants and animals, but importantly, that the effectiveness of selection at removing variation at linked sites is correlated with population size (Corbett-Detig *et al.* 2015). The increased effectiveness of natural selection in larger populations reduces genetic diversity to a much greater extent than in smaller populations, countering to some degree the reduced influence of genetic drift.

However, this correlative evidence fails to capture the complexities of how natural selection acts in different species. A range of factors will affect the efficiency of natural selection, and how strongly it influences linked sites, including the recombinational landscape across the genome, the frequency of adaptive change and historical population demography (Cutter and Payseur 2013). These factors vary enormously between species, and in-depth analyses of an increasing number of taxa has revealed that not all conform to the same general trends. For example, certain plant species do not show a correlation between recombination and neutral polymorphism (see Cutter and Payseur [2013] for a thorough review). In the insects, most of what we have learnt about the action of selection and drift in natural populations comes from studies of the genus *Drosophila* (Andolfatto 2007; Sella *et al.* 2009; Sattath *et al.* 2011; McGaugh *et al.* 2012; Campos *et al.* 2014; Comeron 2014; Lee *et al.* 2014). For example, in *Drosophila simulans*, genetic diversity is strongly reduced in the vicinity of recent non-synonymous substitutions, indicative of strong hitchhiking around beneficial mutations (Sattath *et al.* 2011; Lee *et al.* 2014). This contrasts with a more subtle pattern in humans, where a reduction in diversity around functional substitutions is only detectable after accounting for background selection (Hernandez *et al.* 2011; Enard *et al.* 2014). It remains to be seen whether the rampant selection seen in *Drosophila* spp. is typical of insects.

Here we investigate the action of selection and other evolutionary forces in *Heliconius* butterflies, focusing in particular on *H. melpomene*. This species differs from *D. melanogaster* in a number of ways that might influence patterns of selection across the genome. Populations of *Heliconius* live in tropical rainforests and are characterised by long life-spans and stable populations (Ehrlich and Gilbert 1973). In addition, *H. melpomene* has a similar per base recombination rate to *D. melanogaster*, but more chromosomes (21 compared to 4), potentially allowing higher overall recombination rates. Although the ecology and evolution of this genus has been the subject of much research (reviewed by Merrill et al. [2015]), including recent genomic studies of adaptation and speciation (Arias *et al.* 2012; Nadeau *et al.* 2012, 2013; Supple *et al.* 2013; Martin *et al.* 2013; Kronforst *et al.* 2013), whole-genome studies of selection and within-species genetic diversity have been lacking. Data from *H. melpomene* was included in the recent comparative study of Corbett-Detig et al. (2015). However, only four individuals from a single population were considered. Here we examine in detail the action and influence of natural selection within and between populations and species using whole genome resequencing data from 59 *H. melpomene* individuals and a further 21 samples from 11 related species. We first identify four large but cohesive populations and then explore genetic variation within and between populations and species, describing the footprints of various selective and neutral processes.

## MATERIALS AND METHODS

### Mapping, genotyping, and estimation of error rates

The analysed genome sequences from 80 butterflies included both published and new data. Sample information and accession numbers are given in Table S1. The 58 wild-caught *H. melpomene* samples cover much of the species range and included 13 wing pattern races. We also re-analysed sequence data from a single individual from the inbred *H. melpomene* reference strain (The *Heliconius* Genome Consortium 2012). For sequences generated in this study, methods were as described by Martin et al. (2013). All sequences analysed here consisted of paired-end reads obtained by shotgun sequencing using either Illumina’s Genome Analyzer IIx system or Illumina’s HiSeq 2000 system, according to the manufacturer’s protocol (Illumina Inc.).

Quality-filtered, paired-end sequence reads were mapped to the *H. melpomene* genome scaffolds (version 1.1) (The *Heliconius* Genome Consortium 2012) using Stampy v1.0 (Lunter and Goodson 2011). Genotypes were called using the GATK v2.7 UnifiedGenotyper (DePristo *et al.* 2011). See File S1 for detailed methods. Only “high quality” genotype calls (Phred-scaled mapping quality and genotype quality ≥ 30) were used in downstream analyses. We optimised our genotype calling procedure by examining the total numbers of genotype calls and estimated error rates produced by different pipelines. The rate of false positive heterozygous genotype calls was estimated by analysing a homozygous region in the inbred reference sample. We further examined how error rates change with increasing divergence and different read depths using simulated sequence reads generated using seq-gen (Rambaut and Grass 1997) and ART (Huang *et al.* 2012). See File S1 for further details.

### Analysis of phylogeny and population structure

A maximum-likelihood tree for all 80 samples was generated using only four-fold degenerate sites that had high-quality genotype calls in at least 60 samples, giving an alignment of 1.7 million bases. RAxML (Stamatakis 2006; Ott *et al.* 2007; Stamatakis *et al.* 2008) was used with the GTRGAMMA model, and 100 bootstrap replicates were performed.

We then used two approaches to identify populations that would be considered separately in downstream analyses of diversity: STRUCTURE (Pritchard *et al.* 2000; Falush *et al.* 2003), a model-based clustering method that infers the proportion of each individual’s genotype made up by each of a defined number of clusters; and Principle Components Analysis (PCA), performed using Eigenstrat SmartPCA (Price *et al.* 2006). To minimise the influence of selection, both analyses considered only four-fold degenerate sites. Detailed methods are provided in File S1.

### Site frequency spectra

We generated unfolded site frequency spectra for each *H. melpomene* population by counting the number of derived alleles at biallelic sites. Sites were polarised by comparison with the ‘silvaniform’ clade species: *H. hecale*, *H. ethilla* and *H. pardalinus*. To allow comparison among populations, and account for missing data, each site was randomly down-sampled to the same number of individuals. See File S1 for details.

### Inference of historical population size change using PSMC

To infer changes in ancestral population sizes, we used the pairwise sequentially Markovian coalescent (PSMC) program (Li and Durbin 2011). This method fits a model of fluctuating population size by estimating the distribution of times to most recent common ancestor across a diploid genome. Twelve samples were selected *a-priori* for PSMC analysis. These twelve were chosen because they all had similar sequencing depth, similar numbers of genotyped sites, were all male (homogametic, ZZ), and provided a good representation across the species range. Detailed methods are provided in File S1.

### Window-based population parameters

Various population parameters were calculated for non-overlapping 100 kb windows across the genome. Only windows with a sufficient number of sites genotyped in at least 50% of samples were considered. see File S1 for details. We used 100 kb windows because linkage disequilibrium (LD) tends to break down almost completely within 10 kb, and reaches background levels within 100 kb (Fig. S1), meaning that measures from adjacent windows would be largely free of linkage effects.

Nucleotide diversity (π) and absolute divergence (*d*_*XY*_), were calculated as the average proportion of differences between all pairs of sequences, either within a sample (π) or between two samples (*d*_*XY*_). Sites with missing data were excluded in pair-wise manner to maximise the amount of data being considered. Tajima’s D (Tajima 1989) and *F*_*ST*_ (as in equation 9 of Hudson et al. [1992]) were calculated using the EggLib Python module (De Mita and Siol 2012).

### Estimating the rate of adaptive substitution

We estimated the genome-wide rate of adaptive substitution (α) using Messer and Petrov’s asymptotic method (Messer and Petrov 2013), comparing synonymous and non-synonymous SNPs covering 11,804 polymorphic genes (11,638 autosomal and 166 Z-lined). Polymorphism was measured in the Western population of *H. melpomene*, and divergence was measured between the Western population and *H. erato*. We calculated confidence intervals around the estimated alpha by performing 1000 bootstraps, in each of which 11,804 genes were re-sampled, with replacement. Detailed methods are described in File S1.

### Multiple Regression

We used multiple linear regression to model nucleotide diversity at 4D sites (π_4D_) in 100 kb windows (calculated for each population separately and then averaged). The aim was to assess the influence of selection at linked sites on diversity at neutral sites. Since linked selection is largely modulated by the number of selected sites and the extent of linkage, we included as explanatory variables the local gene density (the proportion of coding sequence per window) as proxy for the density of nearby selected sites (Corbett-Detig *et al.* 2015) and local recombination rate 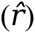, calculated from the linkage map. To account for genetic hitchhiking, we also included the number of recent non-synonymous substitutions (*D*_*n*_) in the *H. melpomene* lineage per window as an explanatory variable. As an alternative, and potentially more direct indicator of adaptive substitutions, we also tested a model using summed gene-by-gene estimates of the number of adaptive non-synonymous substitutions (*a*), estimated by maximum-likelihood using the McDonald-Kreitman test framework (Welch 2006). To account for mutation rate variation, the rate of synonymous substitutions per synonymous site (*d*_*S*_) was also included as an explanatory variable. Lastly, we also included GC-content at third codon positions, to account for any effects of DNA composition. Detailed methods for the estimation of various explanatory variables and data processing for this analysis are provided in Text S1.

To further investigate the interrelationships between the explanatory variables, we used principal component regression (PCR) (Drummond *et al.* 2006; Mugal *et al.* 2013). This approaches can help to tease apart the effects of the various explanatory variables by summarizing the explanatory variables into orthogonal components, thereby accounting for multi-collinearity. Regression analyses were performed with the R version 3.0.3 (https://www.R-project.org) using the *pls* package (Mevik BH, Wehrens R 2013).

To assess the robustness of our findings, we investigated the influence of various modifications to the model, such as restricting the analysis to genes showing minimal codon usage bias, the exclusion of chromosome ends, and use of a different outgroup. Details are provided in Text S1.

Multiple linear regression was also performed for whole chromosomes, where the response variable was the mean 4D site diversity per chromosome 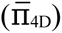. Here, rather than using recombination rate estimated from the linkage map, we used chromosome length as a proxy for recombination rate (Kaback *et al.* 1992; Lander *et al.* 2001). Thus, the five explanatory variables were chromosome length, average gene density, average synonymous substitution rate 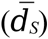, average number of non-synonymous substitutions per 100 kb 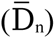 and average GC-content. As above, 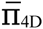 was square-root transformed, but in this model none of the explanatory variables required transformation to correct for skewness. As above, all explanatory variables were Z-transformed so that their effects could be compared.

### Scanning for selective sweeps

To identify candidate selective sweep locations in the Eastern and Western populations, we used SweeD (Pavlidis *et al.* 2013). This program is based on Sweepfinder (Nielsen *et al.* 2005), and uses a composite likelihood ratio (CLR) to identify loci showing a strong deviation in the site frequency spectrum towards rare variants. See Text S1 for detailed methods.

### Data Availability

All raw sequence reads are available from the Sequence Read Archive. Accession numbers are provided in Table S1. Processed genotype data, along with data files underlying all results, figures and tables, and code used for model fitting are available from Data Dryad (http://datadryad.org/review?doi=doi:10.5061/dryad.g0874).

## RESULTS

### Genotyping

The median depth of coverage across all samples was 28x. A median of 78% of sites genome wide, and 96% of coding sites, had high-quality genotype calls for samples of *H. melpomene*, its two close relatives *H. cydno* and *H. timareta*, and the ‘silvaniform’ clade species *H. hecale*, *H. ethilla* and *H. pardalinus*, which diverged from *H. melpomene* about 3.8 million years ago (Ma) (Kozak *et al.* 2015) (Fig. S4). A few samples had considerably fewer sites genotyped, owing to poor sequence coverage (Table S1). More distant species, including *H. wallacei* (~8.8 Ma), *H. doris* (~9.7 Ma) and *H. erato* (~10.5 Ma) all showed strongly reduced numbers of genotype calls (median 33%), suggesting that many reads from these species were too divergent to be mapped reliably to the *H. melpomene* reference. However, the number of calls obtained in coding regions showed very little drop-off with phylogenetic distance, with a median of 90% of coding sites genotyped in the most divergent outgroup, *H. erato* (Fig. S4). This implies that coding regions are sufficiently conserved to allow read mapping and genotyping across all *Heliconius* species. Our analyses of the more distant species therefore focused only on coding regions.

We selected a genotyping pipeline that gave a false positive SNP rate of 0.03% per site (3 errors in 10,000 calls), when comparing the inbred reference sample to itself (Table S2). Using simulated reads, we found that our pipeline produced higher error rates for more divergent taxa, especially when sequencing depths were low (Fig. S5). Nevertheless, for divergences below 6%, which is typical for coding sequences in this genus, and with appreciable sequencing depth, estimated error rates were still well under 0.05% (5 in 10000). As we are concerned primarily with large-scale genomic trends, with all analyses considering large numbers of sites, rare genotyping errors are unlikely to influence our conclusions.

### Population structure and phylogenetic relationships

In order to focus our analyses on biologically meaningful populations, we first investigated population structure among our samples. Analysis of population structure based on four-fold degenerate (4D) sites, using both Principal Components Analysis (PCA) and STRUCTURE (Pritchard *et al.* 2000; Falush *et al.* 2003) gave largely congruent results. Both analyses identified three distinct *H. melpomene* clusters that were largely partitioned geographically (Fig. 1B and C). Consistent with previous studies using smaller datasets, *H. melpomene* samples from the eastern and western slopes of the Andes formed two strongly-differentiated populations, separated by a deep phylogenetic split (Fig. 1B). The third population was made up of the samples from French Guiana. These three populations will be referred to as the “Eastern”, “Western” and “Guianan” populations. The only exception to this geographic clustering was a group of five samples of *H. m. melpomene* from the eastern slopes of the Andes in Colombia, which formed a monophyletic clade most closely allied with the Western population. However, the STRUCTURE results suggested admixture between these and both the Eastern and Guianan populations (Fig. 1B). Although not differentiated by principal components 1 and 2 (Fig. 1C), these five Colombian samples were differentiated from the Western population by principal component 3 (Fig. S9). Given the distinct geography and genomic composition of these samples, we made the conservative decision to consider this group as a fourth distinct population (“Colombian”).

**Fig. 1.**
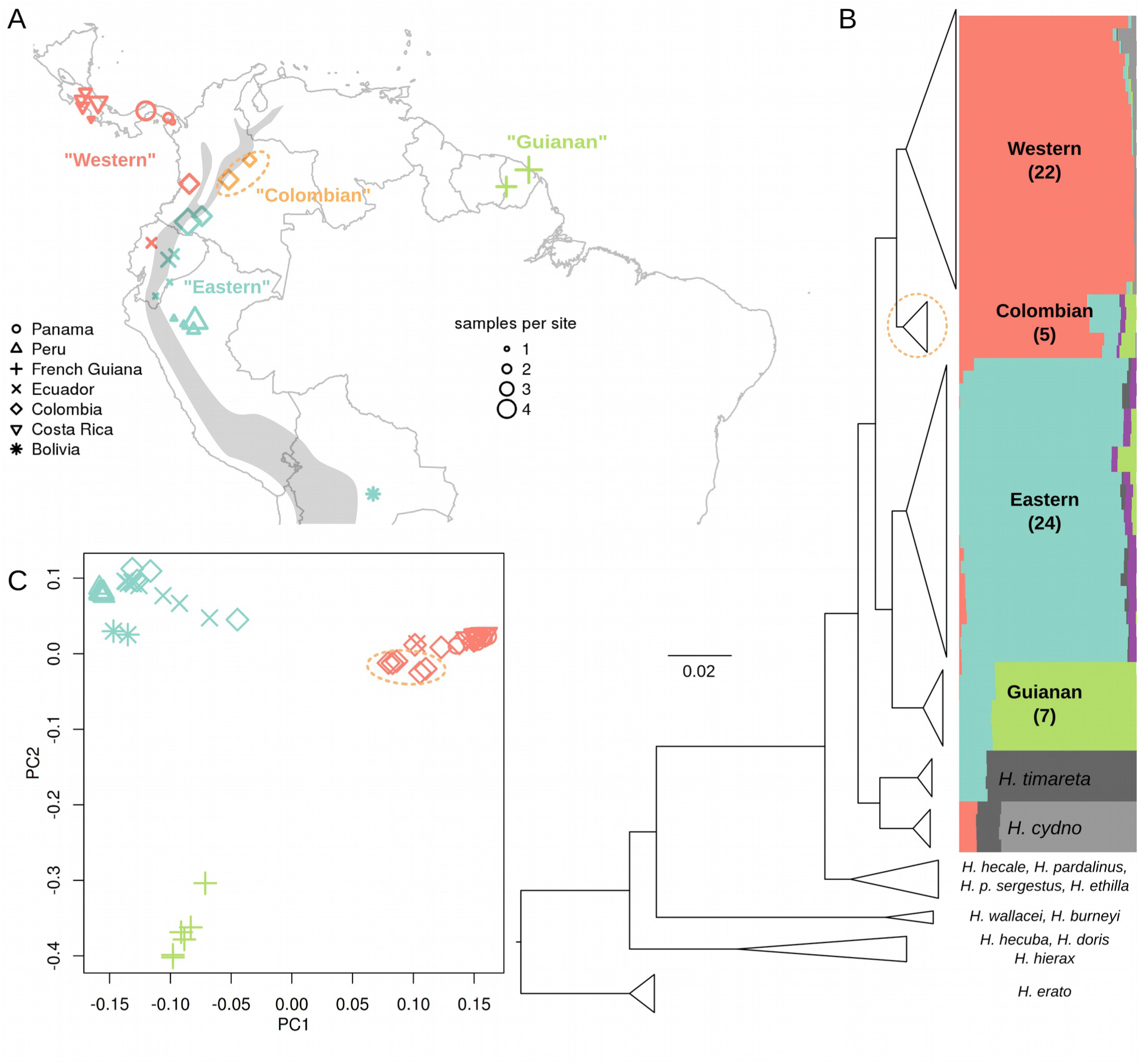
Sample locations, phylogeny and population structure. **A**. Sampling locations of the 58 wild *H. melpomene* samples (see Table S1 for coordinates). Symbols indicate country of sampling, sizes indicate the number of samples from each location. Colours correspond to major clustering on the STRUCTURE plot (B). Grey shading indicates the approximate location of the Andes mountains. **B**. Compressed RaxML phylogeny based on four-fold degenerate (4D) sites. See Fig. S6 for an uncompressed version. Coloured bars indicate genotype cluster proportions for each sample inferred by STRUCTURE with k=6. STRUCTURE plots for k=5-8 are given in Fig. S7, and Ln probabilities for different k values are given in Fig. S8. **C**. Principal Component 1 plotted against Principal Component 2, which explained 17% and 7.7% of the variance, respectively. Colours and symbols as in A. In A, B and C the Colombian samples discussed in the text are circled in orange.

### Recent expansion of the Eastern population

Tests for selection can be confounded by historical changes in population size (Eyre-Walker and Keightley 2009; Li *et al.* 2012), so we investigated the population history of the four *H. melpomene* populations. Three of the four populations showed signatures consistent with fairly stable population sizes, but the Eastern population showed evidence of a recent expansion. Tajima’s D was consistently negative in the Eastern population, but close to zero in Western, Colombian and Guianan populations (Fig. 2A). Negative Tajima’s D is indicative of an excess of rare variants, consistent with recent population growth. This finding was substantiated by direct examination of the unfolded site frequency spectrum (SFS). The SFS at 4D sites showed a strong excess of rare variants in the Eastern population compared to the other populations (Fig. 2B). This skew remained when only geographically proximate samples, with high sequencing coverage were considered (Fig. S10), indicating that it was not an artefact of sampling design. The same trend was also observed at intronic and intergenic sites (Fig. S10). Compared to neutral expectations with constant population size, the other three populations displayed a weak excess of rare variants, but to a much lesser degree than the Eastern population. For the Eastern and Western populations, which were more densely sampled, we were able to compare the SFS down-sampling to 20 individuals for each SNP position. This deeper sampling reproduced the pattern, further showing that the skew was not limited to singleton variants, but also doubletons (derived alleles present twice in the sample) (Fig. S10). While genotyping error could explain some of the excess of singleton SNPs, it is unlikely to cause the observed excess of doubletons, nor the dramatic skew seen in the Eastern population.

**Fig. 2.**
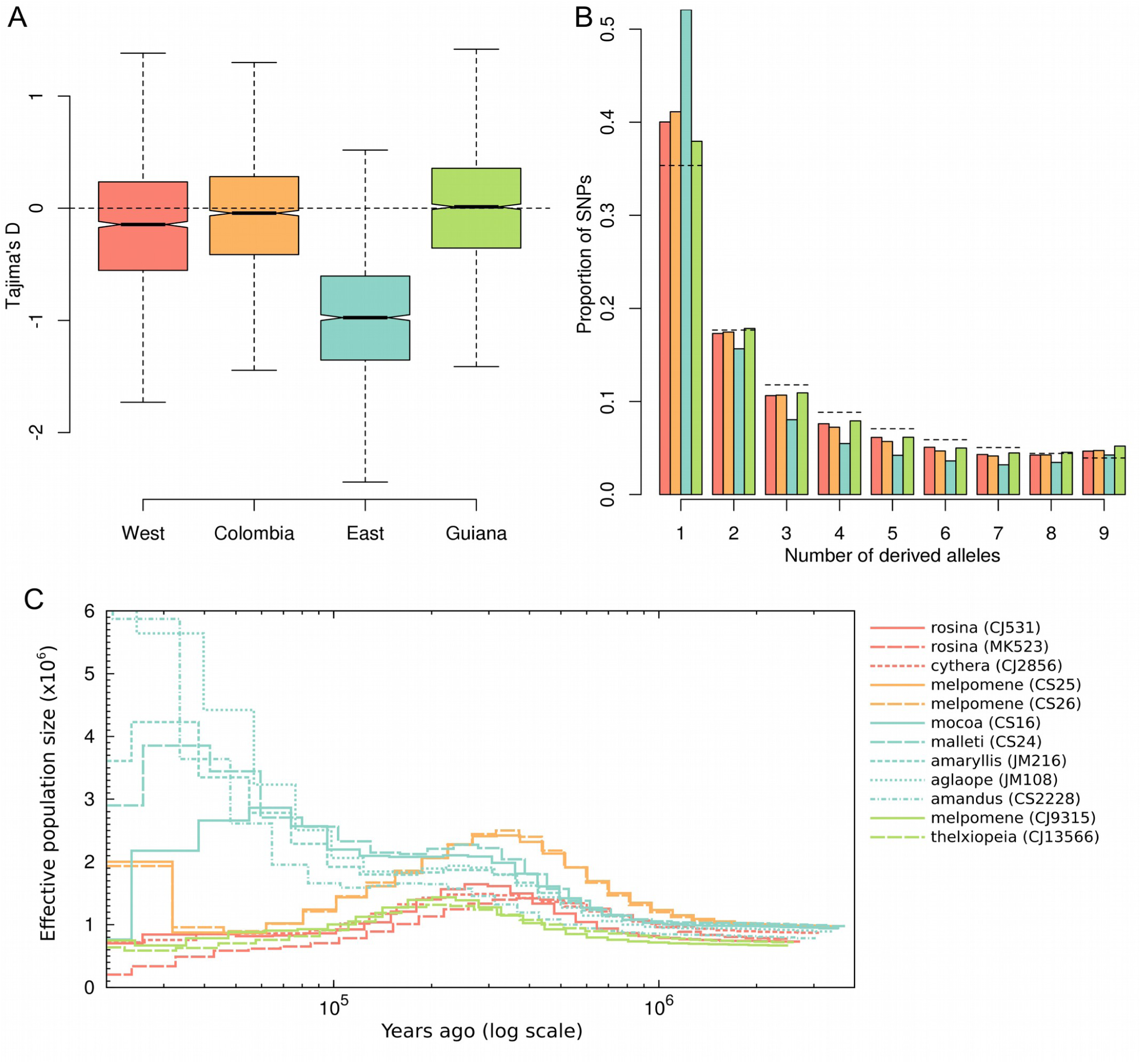
Evidence for recent expansion of the Eastern population. **A**. Boxplots of Tajima’s D, calculated for 4D in each 100 kb windows throughout the genome. **B**. Site frequency spectra for four-fold degenerate (4D) sites for the four *H. melpomene* populations, sampling five individuals per site. Colours denote populations; red: Western, orange: Colombian, blue: Eastern, green: Guianan. Dashed lines indicate the expected frequencies under the standard coalescent model with constant population size (Fu 1995). **C**. PSMC plots of inferred population size through time on a logarithmic axis. Twelve selected male samples that had similar numbers of genotyped sites were included. Source populations are coloured as in A and B.

We further verified our hypothesis of a recent expansion in the Eastern population using the pairwise sequentially Markovian coalescent (PSMC) method of Li and Durbin (Li and Durbin 2011). All four populations showed similar population size histories up until about 200,000 years ago, with a gradual increase in population size beginning about 1 Ma and levelling off around 300,000 years ago. However, while the Western, Colombian and Guianan samples showed a subsequent decrease in the inferred *N*_*e*_, that of the Eastern samples rose again, roughly doubling between 100,000 and 30,000 years ago (Fig. 2C). Closer to the present (< 30,000 years ago) the inferred individual histories diverged considerably, as may be expected given the dearth of information about recent demography to be gained from analysis of single genomes (Li and Durbin 2011). Nevertheless, there appears to be a tendency for the more southern samples to show greater expansion (e.g *H. m. amandus* from Bolivia and *H. m. aglaope* from Peru). Methods more sensitive to demographic change in the recent past may be necessary to confirm this pattern. This analysis was repeated several times, varying the PSMC input parameters for block size and recombination rate. While absolute population size estimates tended to be lower at smaller block sizes, the observed trend of a population size expansion in each of the Eastern samples was consistent throughout (data not shown). As it is based on heterozygosity in single genomes, this analysis is independent of the site frequency spectrum, and therefore provides an additional line of evidence for a recent expansion of the *H. melpomene* population to the east of the Andes.

One potential caveat in this conclusion is that hybridisation and gene flow may also produce patterns consistent with population growth, and gene flow is known to occur between the eastern population and *H. timareta* (Martin *et al.* 2013). However, there is also significant gene flow between the Western population and *H. cydno*, implying that hybridisation alone is unlikely to explain the distinct pattern seen in the Eastern population.

### Diversity and divergence across the genome

Estimated neutral diversity in *H. melpomene* was found to be high, and comparable with that in *Drosophila* spp. Estimates of within-population nucleotide diversity (π) in *H. melpomene* made use of only those samples with at least 25x depth of coverage, because we found that levels of within-sample heterozygosity tended to be underestimated at sequencing depths below this threshold (Fig. S11). Genome-wide π, averaged over all 100 kb windows across the four populations was 1.9%, and similar when only intergenic (2.0%) or intronic (1.9%) sites were considered (Table 1, Table S3). As expected, diversity was strongly reduced at first and second codon positions (0.6%) and higher at third codon positions (1.5%). Diversity was highest at 4D sites (2.5%).

**Table 1.** Nucleotide diversity (π) in *H. melpomene* and absolute divergence (*d*_*XY*_) from outgroups.

| Site class | $\pi$<br>(*melpomene*) | *d<sub>XY</sub>* between *melpomene* and ... | | | | |
| --- | --- | --- | --- | --- | --- | --- |
|  |  | *cydno*,<br>*timareta* | *hecale*,<br>*ethilla*,<br>*pardalinus* | *wallacei*<br>*burneyi* | *doris*,<br>*hecuba*,<br>*hierax* | *erato* |
| Whole genome |  |  |  |  |  |  |
| All sites | 0.019 | 0.027 | 0.036 | . | . | . |
| intergenic | 0.020 | 0.029 | 0.038 | . | . | . |
| intron | 0.019 | 0.028 | 0.038 | . | . | . |
| Codon 1 | 0.006 | 0.010 | 0.014 | 0.026 | 0.022 | 0.037 |
| Codon 2 | 0.006 | 0.009 | 0.012 | 0.022 | 0.020 | 0.032 |
| Codon 3 | 0.015 | 0.024 | 0.033 | 0.065 | 0.056 | 0.091 |
| 4D | 0.025 | 0.041 | 0.057 | 0.114 | 0.100 | 0.158 |
| Z chromosome |  |  |  |  |  |  |
| All sites | 0.011 | 0.024 | 0.034 | . | . | . |
| intergenic | 0.012 | 0.025 | 0.035 | . | . | . |
| intron | 0.011 | 0.025 | 0.036 | . | . | . |
| Codon 1 | 0.003 | 0.008 | 0.013 | 0.028 | 0.024 | 0.036 |
| Codon 2 | 0.003 | 0.007 | 0.012 | 0.023 | 0.021 | 0.030 |
| Codon 3 | 0.007 | 0.019 | 0.030 | 0.064 | 0.059 | 0.094 |
| 4D | 0.011 | 0.032 | 0.049 | 0.107 | 0.100 | 0.158 |

See Table S3 for full data including error margins.

The peripheral Western and Guianan populations had significantly lower diversity than those at the centre of the range (Fig. 3A) (paired Wilcoxon Signed Rank Test, P<2e-16). To ensure that this trend was not simply driven by population sub-structure among sampled individuals, we examined levels of heterozygosity within each sample. As mentioned above, this revealed that sequencing depth affected estimates of heterozygosity, but that above a depth of roughly 25x, heterozygosity was consistent within each population. Considering only samples with depth of at least 25x, we found that average 4D site heterozygosity in the Western samples (2.67%) was only marginally lower than that in the Colombian (2.79%) and Eastern (2.82%) samples, whereas that of the Guianan samples (2.16%) remained considerably lower than the other populations (Fig. S11).

**Fig. 3.**
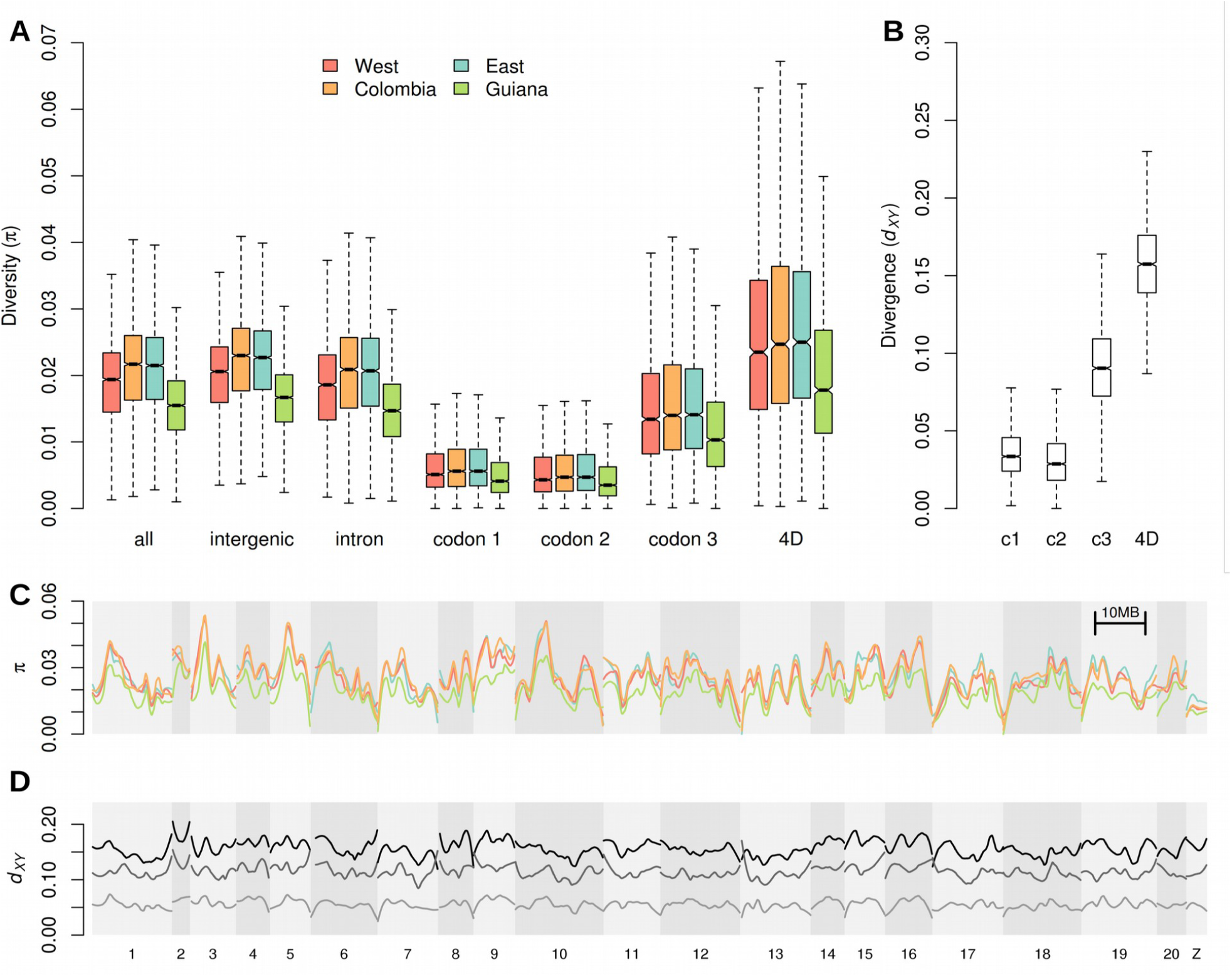
Genome-wide diversity and divergence. **A**. Boxplots of nucleotide diversity (π) for different site classes in the four *H. melpomene* populations. π values were calculated in 100 kb windows, considering all sites of each class within each window. **B**. Boxplots of divergence (*d*_*XY*_) between *H. melpomene* and *H. erato* at four site classes: first second and third codon positions and four-fold degenerate (4D) sites. Note the different y-axis scale **C**. Nucleotide diversity at 4D sites plotted across the 21 *H. melpomene* chromosomes (shaded). Scaffold order was inferred from the *H. melpomene* genome linkage map of v1.1. Populations are coloured as in A. Values are for non-overlapping 100 kb windows, smoothed with loess (local regression), with a span equivalent to 4 Mb. **D**. Divergence (*d*_*XY*_) across the genome, between *H. melpomene* and the silvaniform clade, *H. doris* clade and *H. erato* (respectively, from light to dark), smoothed as in C.

To estimate as closely as possible the value of θ=4*N*_*e*_*μ*, we re-calculated average diversity at 4D sites considering only autosomal genes showing minimal codon usage bias. This gave a slightly higher value of 2.7%, ranging from 2.1% to 2.9% across the four populations (Table 2). These values are in the same range as estimated neutral diversity for *D. melanogaster* in in Southern Africa (~2%) and *Drosophila simulans* (~3.5%) (Begun *et al.* 2007; Langley *et al.* 2012).

**Table 2.** Estimated neutral π (θ) for the four populations, and corresponding population size estimates (in millions) given two different mutation rates.

| Population | Neutral $\pi$ ($\theta$) | $N_e$ ($\times 10^6$) [$\mu = 2.90 \times 10^{-9}$] | $N_e$ ($\times 10^6$) [$\mu = 1.90 \times 10^{-9}$] |
| --- | --- | --- | --- |
| Western | 0.028 | 2.385 | 3.641 |
| Colombia | 0.029 | 2.503 | 3.821 |
| Eastern | 0.029 | 2.469 | 3.769 |
| Guiana | 0.021 | 1.820 | 2.778 |

Mean divergence at 4D sites between *H. melpomene* and *Heliconius erato*, its most distant relative in the genus, was 15.8% (16% when only minimal-CUB genes were considered). A calibrated phylogeny places the split between these two species at approximately 10.5 Ma (Kozak *et al.* 2015). Assuming four generations per year, this corresponds to a neutral mutation rate of 1.9Χ10^−9^ per site per generation. This is about two thirds of the spontaneous mutation rate recently estimated using whole-genome sequencing of parents and offspring in *H. melpomene* (2.9Χ10^−9^) (Keightley *et al.* 2014). Using these two rates, we estimated *N*_*e*_ (θ/4μ) for the four populations, which ranged from 1.8-2.8 million for the Guianan population to 2.5-3.8 million for the Colombian population (Table 2). We note that these do not represent instantaneous values, but rather aggregates over the course of the coalescent time scale. They are also based on a small subset of putatively neutral sites, making them difficult to compare with the PSMC results (Fig. 2C), as the latter are based on the whole genome data, for which the level of polymorphism is around 30% lower (Table 1).

Levels of diversity at 4D sites varied considerably across individual chromosomes (Fig. 3C). This heterogeneity was strongly conserved between the three populations. Inter-specific divergence also varied across the chromosomes, but to a lesser extent (Fig. 3D). Diversity was also reduced on the Z chromosome relative to autosomes, as expected given its lower effective population size (*N*_*e*_). This discrepancy between autosomes and Z was also present in measures of divergence between *H. melpomene* and its closer relatives, but disappeared at higher levels of divergence, with *d*_*XY*_ between *H. melpomene* and *H. erato* being nearly identical for autosomes and Z (Table 1). This is consistent with a decreasing contribution of *N*_*e*_ to coalescence time for deeper species splits.

One potential concern is that highly variable regions may have been missed due to poor read mapping, in which case diversity and divergence might be underestimated. To test for this bias, we investigated whether π and *d*_*XY*_ were correlated with the proportion of missing data per window (measured as sites genotyped in fewer than 50% of samples). Third codon positions averaged just 2.3% missing data among the *H. melpomene* samples, and just 2.4% across the entire set of 79 wild samples. Neither π nor *d*_*XY*_ were correlated with the proportion of missing data (Fig. S12, S13). Hence, there is no evidence for any bias in coding regions. In intergenic regions, which averaged 23% missing data in *H. melpomene* samples and 25% across the whole sample set, both π and *d*_*XY*_ were found to be weakly correlated with the proportion of missing data (Fig. S12). However, the amount of variance explained by missing data was very low (linear regression, R^2^ = 0.037 and 0.009, respectively). The effect of missing data on our estimates of diversity and divergence therefore appears to be minimal. We suggest that the reduced rate of read mapping in non-coding regions in *H. melpomene* and its closer relatives is probably more driven by an abundance of repeats and structural variation, rather than excessively divergent sequences.

### The rate of adaptive fixation

We estimated that 31% of fixed amino acid substitutions between *Heliconius* species are adaptive. We used Messer and Petrov’s asymptotic method (Messer and Petrov 2013) to estimate a genome wide α, the proportion of non-synonymous substitutions driven by positive selection. The exponential model showed a good fit to the data (Fig 4), and gave an estimated α of 31.0%. The 5^th^ and 95^th^ quantiles from 1000 bootstrap replicates were 29.0% and 33.0%, respectively. This value is roughly intermediate between estimates for humans (13%) and *D. melanogaster* (57%), generated using the same approach.

**Fig. 4.**
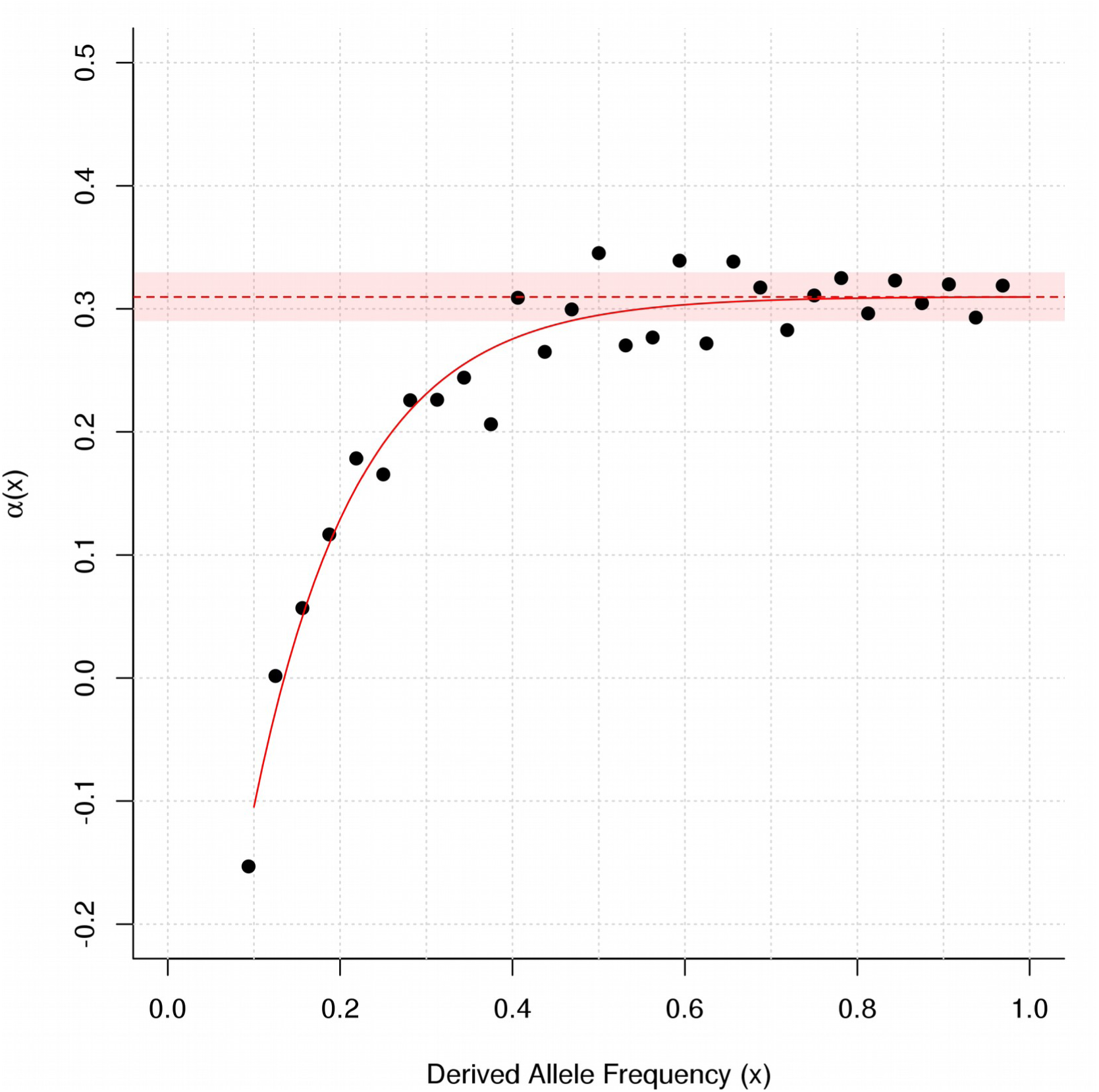
Estimating the rate of adaptive substitution. Mean genome-wide α was estimated using the ‘asymptotic’ method (Messer and Petrov 2013), based on polymorphism for the Western population of *H. melpomene*, and divergence between the Western population and *H. erato*. α(*x*) was calculated for each derived allele frequency (*x*) for all *x* ≥ 0.1. The solid red line indicates the fit of the asymptotic exponential function α(*x*) = *a* + *b*exp(-*cx*), extrapolated to *x*=1 (dashed red line). The solid red box indicates the range between the 5^th^ and 95th percentiles from 1000 bootstrap samples over the 11,804 genes used.

### Selection reduces diversity at linked neutral sites

We explored the influence of selection on linked sites using a multiple linear regression approach, following several previous studies (Cutter and Moses 2011; McGaugh *et al.* 2012; Mugal *et al.* 2013). Our ‘main model’ had nucleotide diversity at 4D sites (π_4D_) in 100 kb windows as the response variable and five explanatory variables: (i) local gene density, a proxy for the number of nearby selected sites; (ii) local recombination rate 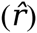; (iii) the number of recent non-synonymous substitutions (*D*_*n*_), to account for hitchhiking around beneficial mutations; (iv) the synonymous substitution rate (*d*_*S*_), a proxy for local mutation rate; and (v) GC-content, to account for effects of local DNA composition (Fig. S14, S15). The results for the main model are summarised in Table 3 and Table S4 and described below. There was limited serial correlation among windows. The Durbin-Watson statistic (Durbin and Watson 1950, 1951) for the main model was 1.3, suggesting that autocorrelation is unlikely to influence our conclusions (Field 2009). Several modifications of the model were also tested to investigate the robustness of the result. These are all summarised in Tables S5-S8. Overall, results were consistent throughout, but several notable differences are described below.

**Table 3.** Summary of multiple regression with five explanatory variables for 4D site diversity, calculated for 100 kb windows (R^2^ = 0.340; adjusted R^2^ = 0.339; F_5,1461_ = 151.1; p < 2.2e-16).

| | Estimate | Std. error | SS | RSS | $F_{1,1461}$ | P (>F) | Partial $R^2$ | VIF |
| --- | --- | --- | --- | --- | --- | --- | --- | --- |
| Gene den. | -0.0127 | 0.0010 | 122.876 | 1230.8 | 162.028 | **< 2.2e-16\*** | 0.0998 | 1.483 |
| $\hat{r}$ | 0.0062 | 0.0008 | 49.496 | 1157.5 | 65.267 | **1.357e-15\*** | 0.0428 | 1.011 |
| $D_n$ | -0.0038 | 0.0010 | 11.432 | 1119.4 | 15.075 | **0.0001\*** | 0.0102 | 1.567 |
| $d_s$ | 0.0147 | 0.0010 | 197.848 | 1305.8 | 260.889 | **< 2.2e-16\*** | 0.1515 | 1.156 |
| GC cont. | -0.0036 | 0.0008 | 14.391 | 1122.4 | 18.977 | **1.416e-05\*** | 0.0128 | 1.076 |
| (Intercept) | 0.1516 | 0.0009 |  |  |  |  |  |  |

$\hat{r}$ : recombination rate, $D_n$ : number of non-synonymous substitutions, $d_s$ : synonymous substitutions per synonymous site, SS: Sum of Squares, RSS: Residual Sum of Squares, VIF: variance inflation factor, \*: $p \leq 0.05$

The main model explained 34% of the variation in π_4D_ (F_5,1461_ = 151.1, p < 2.2e-16) and all five predictor variables were found to have significant effects (Table 3). Unsurprisingly, synonymous substitution rate (*d*_*S*_) was a strong predictor of intra-specific diversity (F_1,1461_ = 260.89, p < 2.2e-16), implying that at least some of the observed heterogeneity in diversity across the genome is explained by variation in mutation rate. Local gene density also showed a strong negative relationship with π_4D_ (F_1,1461_ = 162.03, p < 2.2e-16), consistent with a considerable effect of selection at linked sites. Local recombination rate showed a positive relationship with diversity (F_1,1461_ = 65.27, p < 1.357e-15), also as expected under linked selection. In addition, recombination rate was not correlated with *d*_*S*_ (Pearson’s R = −0.01, p < 0.648, Fig. S16), implying that the positive relationship with diversity cannot be explained by a mutagenic effect of recombination. We suggest that the true relationship between recombination rate and neutral diversity may be stronger than our model predicts, as the imperfect placement of scaffolds on the Hmel1.1 linkage map limits our ability to accurately estimate local recombination rates (see below).

### Evidence for genetic hitchhiking

There was also a significant negative effect of the number of non-synonymous substitutions per window (*D*_*n*_) on π_4D_ (F = 15.08, p < 0.0001) implying a small but detectable effect of genetic hitchhiking around adaptive substitutions. While multi-collinearity among the explanatory variables was generally weak, there was a moderate correlation between gene density and *D*_*n*_ (Pearson’s R = 0.53, Fig. S16). This is unsurprising, since non-synonymous substitutions can occur only in coding sequence, and are thus likely to be more common in gene-rich regions. This can make it difficult to separate the roles of hitchhiking and background selection. However, the variance inflation factors for gene density and *D*_*n*_ in our model were low and do not suggest a significant impact of collinearity on the precision of our estimates (Table 3). Moreover, principal component regression (PCR) analysis, which first accounts for collinearity among explanatory variables by separating them into principle components before performing a multiple regression, demonstrated an effect of gene density on diversity independent of *D*_*n*_ (Fig. S17).

One potential concern is that *D*_*n*_ could be underestimated in highly divergent genes, where read mapping for the distant outgroup *H. erato* may be poor. We therefore tested a modified model in which *d*_*S*_ and *D*_*n*_ were estimated using only the more closely related silvaniform species as outgroups. This made little difference to the results (Table S5).

As an alternative to *D*_*n*_, we also tested a modified model that included maximum-likelihood estimates of *a* (the number of adaptive non-synonynmous substitutions) per gene, summed across each window. This revealed a similar significant negative effect of *a* on π_4D_ (Table S6), while collinearity with gene density was considerably lower than for *D*_*n*_ (Pearson’s R = 0.27). Taken together, these finding all support a role for genetic hitchhiking in addition to background selection in shaping neutral variation in *H. melpomene*.

One striking difference between humans and fruit flies is that genetic hitchhiking in *Drosophila* spp. is pervasive enough to produce an average trough in diversity around non-synonymous substitutions (after scaling for mutation rate variation) (Sattath *et al.* 2011; McGaugh *et al.* 2012); whereas this is not directly observable in humans (Hernandez *et al.* 2011). We performed an equivalent test with our data (Fig. S18), and found patterns similar to those in humans, with no significant reduction of scaled diversity in the vicinity of non-synonymous substitutions compared to synonymous substitutions. Enard et al (2014) show that the effect of hitchhiking around non-synonymous substitutions can be masked by background selection. This may be because background selection tends to be stronger in more conserved genomic regions, where adaptive substitutions are expected to be less common. This might explain the fact that we only observe evidence for reduced diversity around non-synonymous substitutions in our multiple-regression model. Although our approach does not model the action of background selection explicitly, by including gene density and recombination rate as explanatory variables, it appears to account to some extent for the confounding influence of background selection. The relative roles of hitchhiking and background selection may be further resolved in the future by explicitly modelling these different processes (Corbett-Detig *et al.* 2015).

### GC content and codon usage bias

In the main model, GC content was found to be negatively correlated with π_4D_ (F_1,1461_ = 18.98, p = 1.416e-05). This may reflect codon usage bias (CUB), with a preference for codons ending in C or G, which would lead to elevated GC-content at genes under stronger selection for codon usage. Indeed, in our analysis of codon usage, genes with a higher GC content at the third codon position tended to display stronger evidence for CUB (Fig. S5). In a modified model where π_4D_ was calculated using only our defined set of minimal-CUB genes, GC content became a non-significant predictor of diversity, whereas effect sizes for all other explanatory variables were similar (Table S7).

### Effect of chromosome ends

We last confirmed that the observed patterns are not predominantly driven by chromosome ends. A model excluding windows in the outer 5% of chromosomes was similar to the main model (Table S8). Generally, effect sizes and p-values were lower, but this may partly reflect the 10% reduction in the number of observations. Interestingly, GC-content was no longer a significant predictor of diversity. This might imply that patterns of codon usage change toward the chromosome ends, but this will require further investigation.

### The relationship between gene density and neutral diversity is clearly visible

Given the large effect of gene density on diversity at 4D sites, we further explored this relationship visually (Fig. 5). There was a clear trend of lower π_4D_ in gene-rich regions, but also a conspicuous increase in variance in regions of lower gene density (Fig. 5A). This is most likely caused by the smaller number of 4D sites available in such regions, resulting in less data and therefore increased noise. We were able to account for this issue in our multiple linear regression model by weighting residuals according to the number of data points available per window, and overall the model showed minimal violation of assumptions (Fig. S14, Fig. S15). On several chromosomes the correlation between gene density and π_4D_ was remarkably clear (Fig. 5B).

**Fig. 5.**
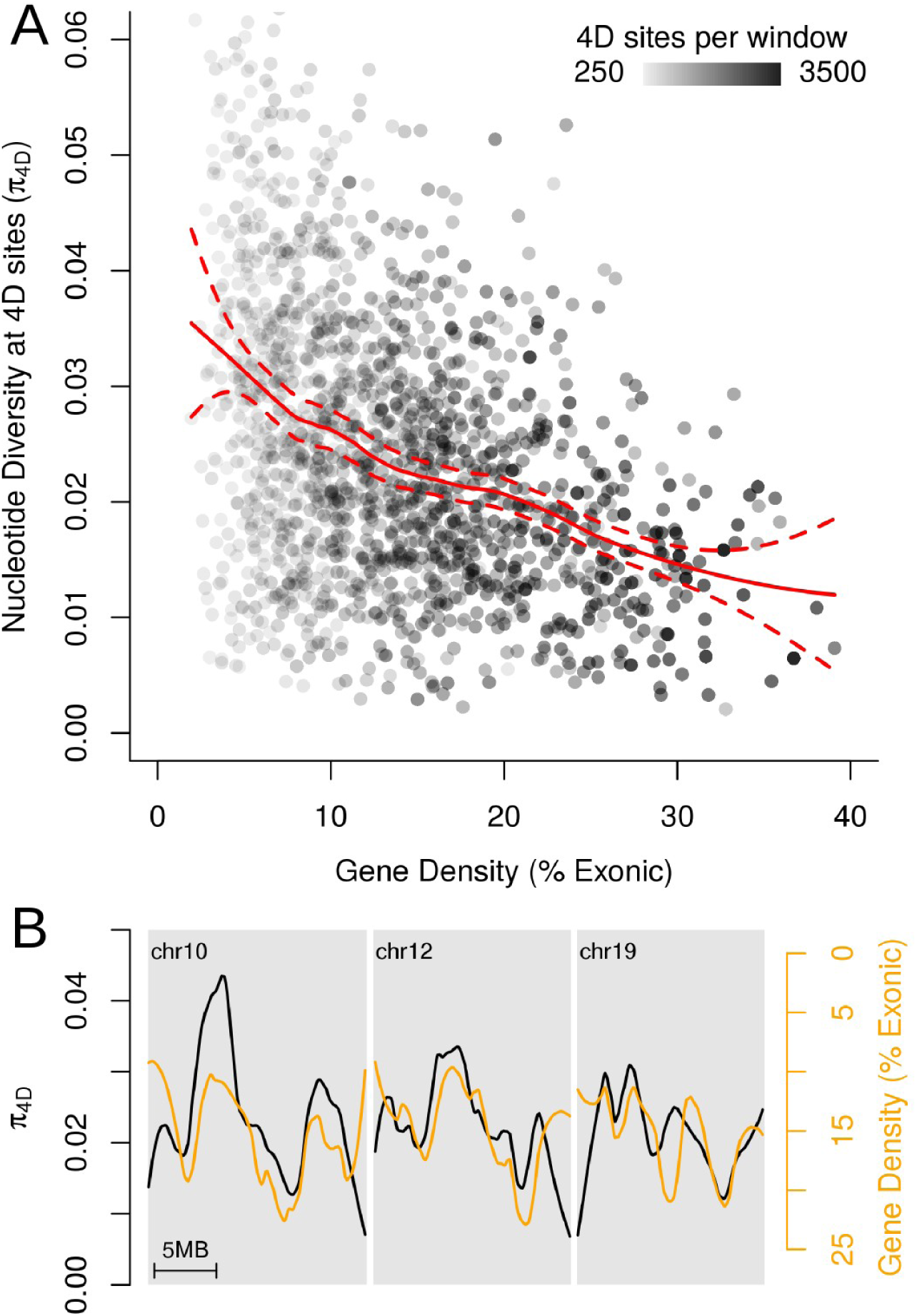
Relationship between gene density and diversity at 4D sites. **A**. Diversity at 4D sites (π_4D_) for 100 kb windows plotted against local gene density, calculated as the percentage of the window made up of exons. Points are shaded according to the number of 4D sites in the window that had genotype calls for at least 50% of analysed samples. A loess (locally weighted smoothing, span=0.5) curve with 99% confidence intervals is shown in red,. **B**. Plots of π_4D_ (black) and gene density (orange) across chromosomes 10, 12 and 19, which all showed a visually striking correlation. Note that the gene-density axis is inverted and adjusted to aid comparison between the lines. Both lines are loess-smoothed with a span equivalent to 4 Mb.

### Longer chromosomes are less polymorphic

There was a strong negative relationship between 4D site diversity and chromosome length (in bases) (Table 4), further supporting the pervasive role of linked selection in shaping genetic diversity in *H. melpomene*. Long chromosomes tend to have lower recombination rates per base pair (Kaback *et al.* 1992; Lander *et al.* 2001; Kawakami *et al.* 2014), which should lead to stronger linked selection. Although a considerable number of scaffolds in the *H. melpomene* v1.1 genome are not properly positioned and oriented on a chromosome, the chromosomal assignment could be inferred for almost all large scaffolds (83% of the genome in terms of bases) (The *Heliconius* Genome Consortium 2012), making for fairly robust estimates of chromosome length. We used a multiple regression model similar to that used for 100 kb windows above, but here averaging all parameters over each of the 20 autosomes, and with chromosome length used as a proxy for recombination rate. This model explained 73.34 % of the variation in average chromosomal diversity at 4D sites, 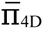 (F_5,14_ = 7.7, p = 0.0011) (Table 4). There was a strong negative relationship between 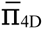 and chromosome length. Comparing models with and without chromosome length as an explanatory variable, we found that the model including chromosome length had a far better fit to the data (F_1,15_ = 20.15, p < 0.0005). Hence, long chromosomes tend to be less variable at neutral sites than short chromosomes, and this trend was clear upon visual inspection (Fig. 6A). As in the window-based model, 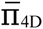 was positively correlated with average synonymous substitution rate, 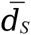(F_1,14_ = 9.85, p = 0.0073), but there was no significant relationship between 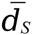 and chromosome length (Pearson’s r = 0.08, p = 0.7289) (Fig. 6B), reinforcing our finding that mutation rates are not correlated with recombination rate. The simplest explanation for this pattern is therefore that linked selection drives patterns of diversity not only among small windows, but also among whole chromosomes.

**Table 4.** Summary of multiple regression with five explanatory variables for mean 4D site diversity calculated for chromosomes (R^2^ = 0.7334; adjusted R^2^ = 0.6382; F_5,14_ = 7.704; p < 0.001144)

| | Estimate | Std. error | SS | RSS | $F_{1,14}$ | P (>F) | Partial $R^2$ | VIF |
| --- | --- | --- | --- | --- | --- | --- | --- | --- |
| Gene den. | 0.0001 | 0.0007 | 1.960e-07 | 8.020e-05 | 0.034 | 0.8557 | 0.0024 | 1.425 |
| Chrom. len. | -0.0034 | 0.0008 | 1.152e-04 | 1.952e-04 | 20.153 | **0.0005\*** | 0.5901 | 1.920 |
| $\bar{D}_n$ | -0.0032 | 0.0013 | 3.318e-05 | 1.132e-04 | 5.807 | **0.0303\*** | 0.2932 | 5.990 |
| $\bar{d}_s$ | 0.0036 | 0.0012 | 5.628e-05 | 1.363e-04 | 9.849 | **0.0073\*** | 0.4130 | 4.495 |
| GC cont. | -0.0019 | 0.0011 | 1.815e-05 | 9.815e-05 | 3.176 | 0.0964 | 0.1849 | 3.722 |
| (Intercept) | 0.0257 | 0.0005 |  |  |  |  |  |  |

$\bar{D}_n$ : average number of non-synonymous substitutions per 100 kb, $\bar{d}_s$ : average synonymous substitutions per synonymous site, SS: Sum of Squares, RSS: Residual Sum of Squares, VIF: variance inflation factor, \*: $p \leq 0.05$

**Fig. 6.**
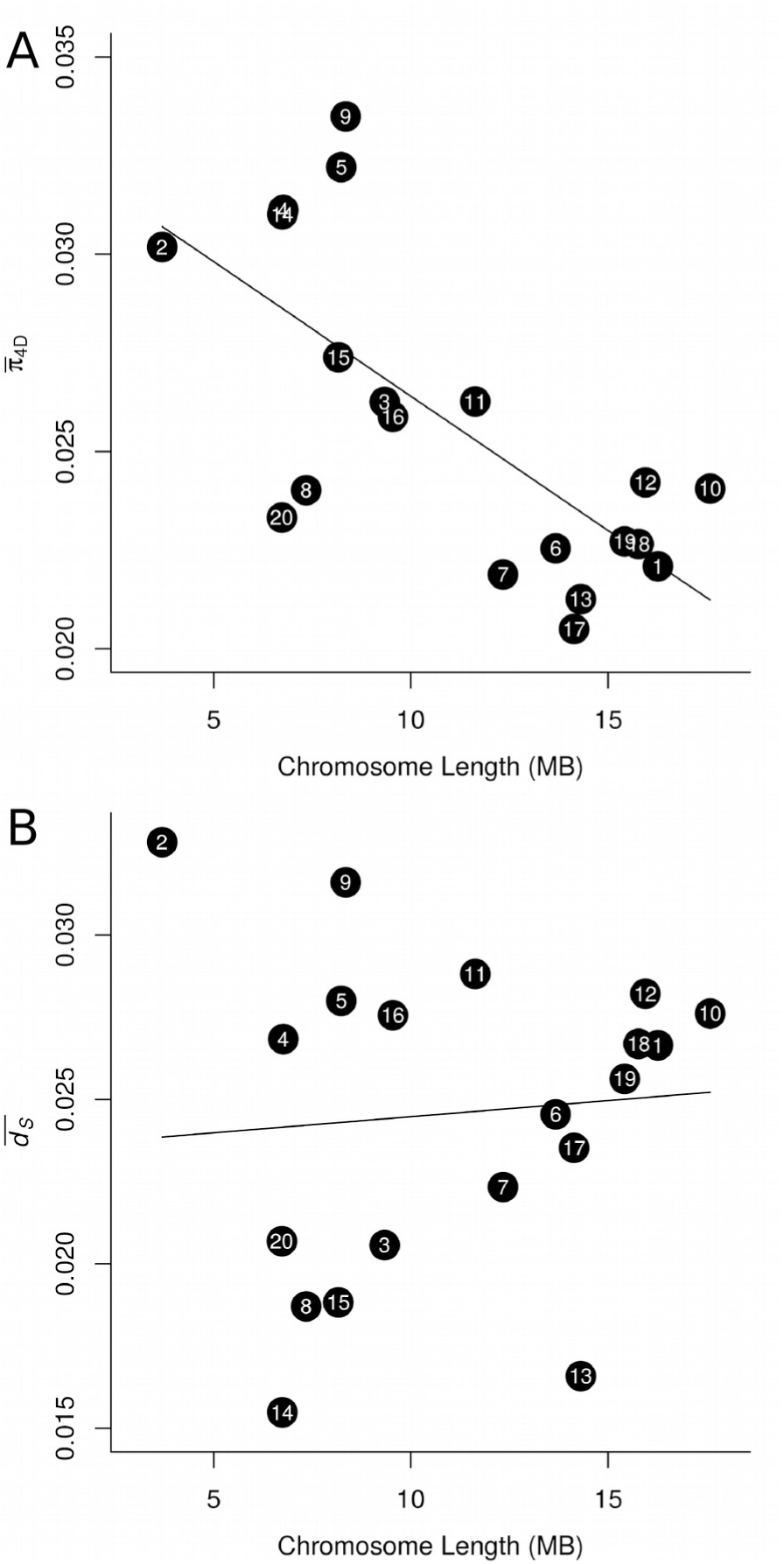
Relationships between chromosome size, 4D site diversity and synonymous substitution rate. **A**. Average 4D site diversity per chromosome 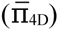 plotted against chromosome length. **B**. The average rate of synonymous substitution per synonymous site per 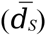 plotted against chromosome length. In both plots, chromosome numbers are indicated, and a linear model fit is shown for reference.

The only other noteworthy correlation with chromosome length was a negative relationship with GC content. We hypothesized that this skew in base composition may be driven by stronger codon usage bias (CUB) on shorter chromosomes. This would be expected if higher recombination rates on shorter chromosomes allowed for more efficient selection by reducing interference among sites (Hill and Robertson 1966; Felsenstein 1974). Consistent with this hypothesis, the proportion of genes identified as having non-negligible CUB was negatively correlated with chromosome length (Spearman’s r = −0.555, df = 19, p = 0.009) (Fig. S19).

### Geographically restricted selective sweeps

We investigated whether there were signatures of strong, recent selective sweeps in *H. melpomene* and also whether sweeps tended to be geographically restricted to a particular population. We scanned the genome for putative selective sweep signals using SweeD (Pavlidis *et al.* 2013), which uses the composite likelihood ratio (CLR) method of Nielsen et al. (2005) to identify loci displaying a strong skew in the site frequency spectrum (SFS) toward rare variants in comparison with the genomic background. A number of regions throughout the genome had outlying CLR values, above a threshold determined using neutral simulations (Fig. S20). Most of these were restricted to the Eastern population, with only one being restricted to the Western population, and one region with partially overlapping outliers in both populations. Given the excess of outliers in the Eastern population, most of which are represented by just single windows, it seems likely that most of these signals are spurious, perhaps reflecting increased variance in the SFS caused by a less stable demographic history. Only two loci had strong putative sweeps indicated by clusters of outlying CLR values. On chromosome 11, outliers in the Eastern and Western populations roughly overlapped (Fig. 7A), consistent with a beneficial allele spreading across both populations. On chromosome 12, the cluster of outlying CLR values were restricted to the Eastern population (Fig. S20). We investigated patterns of diversity within and divergence between populations at these two loci in finer detail. The results were very similar for both candidate sweep loci, so we present the results for chromosome 11 in Fig. 7, and those for chromosome 12 in supplementary Fig. S21.

**Fig. 7.**
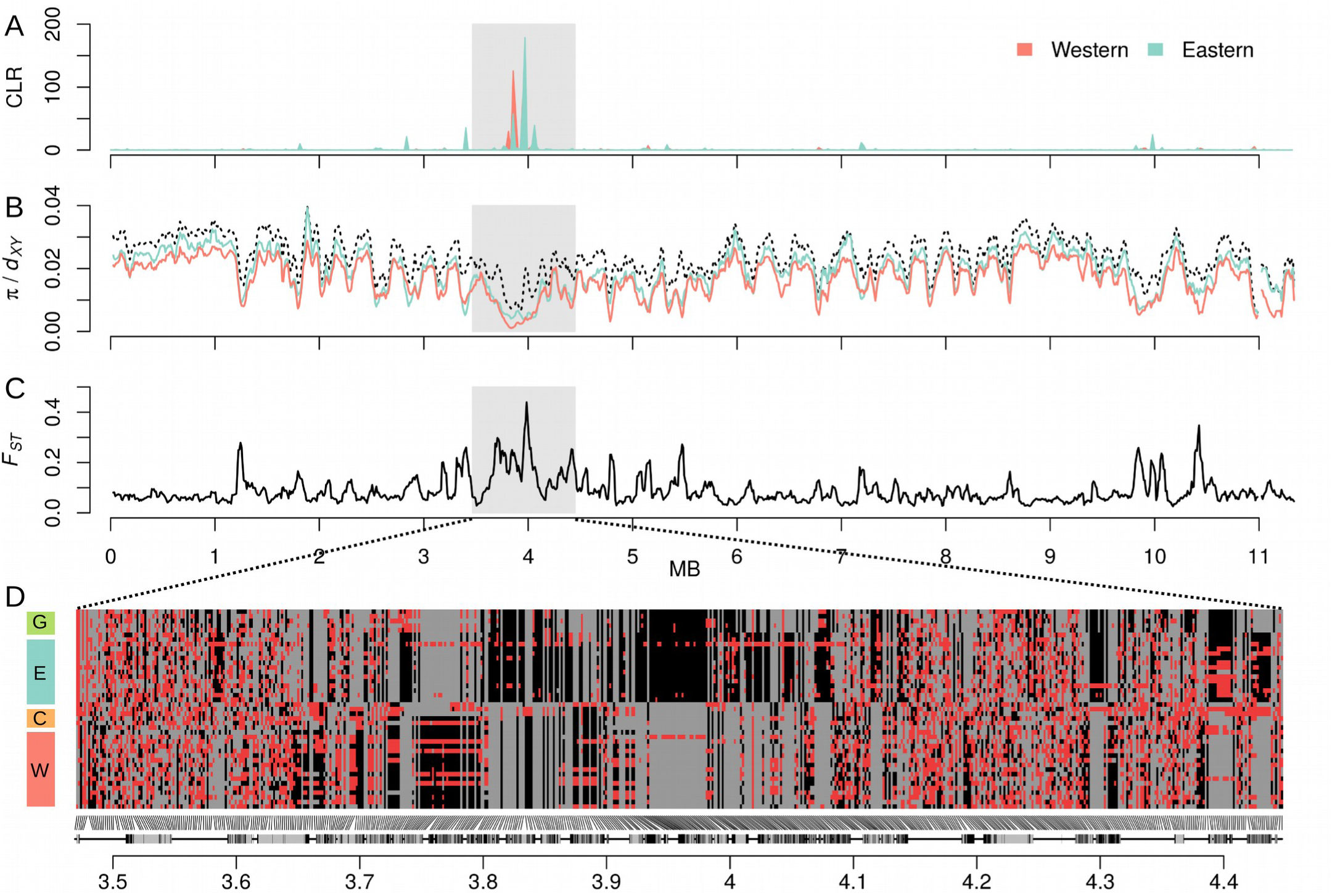
A putative selective sweep on Chromosome 11. A. Composite likelihood ratio (CLR) values calculated by SweeD (Pavlidis *et al.* 2013) for the Eastern and Western populations for 1000 windows across chromosome 11. Scaffolds are shaded light and dark. B. Nucleotide diversity (π) for the Eastern and Western populations (in colour) and divergence (*d*_*XY*_) between these two populations (black dashed line), calculated for 50 kb sliding windows across chromosome 11, sliding in increments of 10 kb. C. FST between the Eastern and Western populations, calculated in windows as in B. D. Individual genotypes at 600 biallelic SNPs on scaffold HE672079, which harbours the putative selective sweep. Homozygous genotypes are coloured grey (major allele) and black (minor allele), and heterozygotes are coloured red. To optimize the detection of differences between populations, SNPs with a high degree of polymorphism (minor allele frequency ≥ 0.25) were considered. The 600 SNPs plotted were sampled semi-randomly, ensuring that no two sampled SNPs were more than 1000 bp apart. Protein coding genes are indicated below the plot, with exons shown in black.

At the putative sweep locus on chromosome 11, nucleotide diversity (π) in both the Eastern and Western populations was reduced (Fig. 7B). Absolute divergence between the two populations (*d*_*XY*_) was similarly reduced (Fig. 7B). However, the fixation index, *F*_*ST*_, between the Eastern and Western populations was strongly elevated in this region (Fig. 7C). This implies that the region carries a low overall amount of genetic variation, but that the variation that is present constitutes fixed or nearly fixed differences between the two populations (Charlesworth 1998). This is consistent with a scenario where the alleles that swept to high frequency in the two populations were not identical, but similar, and potentially carried the same beneficial allele (Bierne 2010). The putative sweep locus on chromosome 12 showed very much the same pattern (Fig. S21).

To further explore the genetic make-up of these regions, we visualised the genotypes of individuals from all four populations at a sample of 600 highly polymorphic bi-allelic SNPs across the scaffold containing the putative sweep locus. Both scaffolds had regions in which heterozygous genotypes were clearly reduced, and fixed differences between the Eastern and Western populations strongly increased. In both cases there were also several fixed differences between the Eastern and Guianan populations, but no fixed differences between the Western and Colombian populations among the 600 sampled SNPs. We note that by focussing on highly polymorphic SNPs, we highlight differentiation between the populations, and fail to show how much of the region is shared, which must be considerable given the reduced *d*_*XY*_. Nevertheless, this visualisation confirms our hypothesis that distinct alleles reached high frequency in the different populations. The presence of some heterozygous sites in the sweep regions indicates that these may be a fairly ancient sweeps and/or that no single allele fixed in each population (i.e. a ‘soft sweep’). One notable observation is the presence of long runs of heterozygous genotypes in certain individuals. This is also consistent with a soft sweep, but may alternatively reflect gene flow subsequent to the sweep, leading to long haplotypes introgressing between the populations.

Both putative sweep locations contained multiple annotated genes. All genes in these two regions that gave significant BLAST hits to *D. melanogaster* proteins (18 and 13 genes, respectively) are listed in Table S9. Due to the large number of potential targets of selection, we do not speculate here as to the adaptive significance of these events.

## DISCUSSION

The extent to which evolutionary change is a result of neutral or selective forces remains one of the long-standing questions in evolutionary biology. Genomic data permit powerful tests of the various forces that shape genetic variation within and between species, but such studies have only recently been extended beyond a few well-studied taxa. We examined a large number of whole genome sequences to investigate the forces shaping diversity in *Heliconius* butterflies. Levels of neutral diversity in *H. melpomene* are similar to those in Southern African populations of *D. melanogaster*, suggesting comparable effective population sizes. However, actual census population sizes of fruit flies must at times reach numbers far greater than those of *Heliconius* butterflies, which are characterised by low-density, stable populations (Ehrlich and Gilbert 1973). This paradox of divergent demography but similar diversity is partly explained by the impact of selection on linked sites. Rampant selection across the compact *Drosophila* genome leads to a dramatic reduction in diversity at linked sites. By contrast, our results suggest that selection is less pervasive in *Heliconius*, and its influence on linked sites, though significant, is less pronounced.

Our findings suggest that positive selection plays an important, but less prominent role in *Heliconius*. We estimate that 31% of amino acid substitutions between *H. melpomene* and *H. erato* were driven by positive selection. This contrasts with an estimated rate of adaptive substitution in *D. melanogaster* of 57%, made using the same method (Messer and Petrov 2013). Although our multiple-regression model indicated that genetic hitchhiking has had a significant effect on neutral variation, we did not detect the same strong reduction in diversity around non-synonymous substitutions seen in *Drosophila* spp. (Sattath *et al.* 2011; McGaugh *et al.* 2012). One possible explanation is that positive selection more often targets non-coding regulatory changes, as appears to be the case in humans (Enard *et al.* 2014). However, even more general signatures of recent selective sweeps in the form of strong skews in the site frequency spectrum were limited. It is important to note that the amount of adaptive evolution depends not just on the efficacy of selection, but also on how often novel, adaptive phenotypes arise. This depends both on the changeability of the fitness landscape and the availability of adaptive variation. Populations of fruit flies may occasionally reach extremely high densities, increasing the potential for selection to detect advantageous mutations (Barton 2010). The relevant population size for adaptive evolution could therefore be much higher than that estimated from neutral variation, especially in species with highly fluctuating populations.

Certain aspects of *Heliconius* biology might also obscure the footprints of positive selection when it does occur. Barriers to dispersal, such as the Andes mountains, can reduce the signature of hitchhiking by slowing the progression of sweeps (Barton 2000; Kim 2013). Indeed, at the two putative sweep loci investigated, similar but distinct alleles appear to have swept in the Eastern and Western populations. This is consistent with a scenario where a globally beneficial allele recombined onto a different genetic background as it spread, which would not only soften the sweep signal, but also enhance population differentiation (Slatkin and Wiehe 1998; Bierne 2010). The source of beneficial variation is another important factor, as adaptation from standing or introgressed variation, can result in soft sweeps (Pennings and Hermisson 2006). One likely example is the repeated evolution of certain wing pattern forms. Despite the strong selection known to act upon wing-patterning loci (Mallet and Barton 1989), signatures of selective sweeps at patter loci have not been observed (Baxter *et al.* 2010; Nadeau *et al.* 2012). While the present study was not designed to address this question, all Western population samples shared a red forewing band, controlled by the *B* locus on chromosome 18 (Baxter *et al.* 2010); and yet no significant sweep signal was detected at this locus. It appears that wing patterning frequently evolves by sharing of pre-existing alleles between populations, and even between species through rare hybridisation (Pardo-Diaz *et al.* 2012; The *Heliconius* Genome Consortium 2012; Wallbank *et al.* 2016). The presence of variation among these old alleles might eliminate any signature of genetic hitchhiking (Pennings and Hermisson 2006). It is yet to be established whether adaptation from standing and introgressed variation is generally common in *Heliconius*, but studies in other systems are increasingly suggesting an important role for pre-existing adaptive variation in evolution (Jones *et al.* 2012; Roesti *et al.* 2014; Gosset *et al.* 2014).

Despite the limited influence of hard selective sweeps, the genomic landscape of variation in *H. melpomene* has been shaped significantly by selection on linked sites. Diversity at neutral sites is positively correlated with recombination rate and negatively correlated with local gene density, which is a good proxy for the density of both coding and non-coding functional elements. These trends are consistent with a pervasive influence of selection on linked sites, and are similar to patterns in several other animals (Cutter and Payseur 2013; Corbett-Detig *et al.* 2015). There is no evidence that recombination rate affects the mutation rate, in agreement with findings in *Drosophila* (McGaugh *et al.* 2012) and humans (McVicker *et al.* 2009). The effects of linked selection are also visible at the whole-chromosome scale. Longer chromosomes are less polymorphic, presumably because they have lower recombination rates per base-pair, leading to stronger effects of linked selection on average. A negative relationship between chromosome length and recombination rate has been observed in a range of taxa (Kaback *et al.* 1992; Lander *et al.* 2001; Kawakami *et al.* 2014), and probably stems from a requirement for at least one obligate crossover event per chromosome during meiosis, even on the smallest chromosomes (Kawakami *et al.* 2014). Increased recombination rates on smaller chromosomes would also be expected to lead to more efficient purifying selection, due to reduced interference among loci (Hill and Robertson 1966; Felsenstein 1974). Indeed, smaller chromosomes display greater evidence of codon usage bias, a phenomenon probably driven by fairly weak selection. This is akin to the observation in *D. melanogaster* of reduced codon usage bias in regions of minimal recombination (Kliman and Hey 1993).

Further studies of other taxa are necessary to build a complete picture of how selection shapes genetic variation in natural populations. Nevertheless, even a simple comparison between butterflies and fruit flies can be enlightening. The different biology of these two insect groups results in distinct patterns of adaptive evolution. However, the lower effective population sizes in *Heliconius* combined with less influence of selection on linked sites leads to levels of neutral diversity similar to those in *Drosophila* spp. Indeed, Corbett-Detig et al. (Corbett-Detig *et al.* 2015) estimate that the impact of selection on linked sites is around three times greater in fruit flies. Although it has long been recognised that levels of neutral variation are not solely determined by population size, whole genome studies such as this are beginning to reveal in detail how different processes combine to shape genetic diversity.

## ACKNOWLEDGEMENTS

We thank the two anonymous reviewers for their comprehensive comments, which greatly improved this manuscript. Nick Barton and Aylwyn Scally provided helpful comments on an earlier draft. We thank John Davey for advice on the recombination rate analyses, and for useful discussions of the results. James Walters provided advice for the analysis of the rate of adaptive substitution, and some of the groundwork for this analysis was performed by Gabriel Jamie and Joseph Harvey. We are also grateful to James Mallet and Lawrence Gilbert for thought-provoking discussions of *Heliconius* life history. Finally, we thank Jenny Barna for computing support. This work was funded by ERC grants 281668 to FMJ and SpeciationGenetics to CDJ. Some sequencing was funded by a John Fell Fund of Oxford University grant to Judith Mank and James Walters. SHM was supported by St John’s College, Cambridge. MM was supported by a SNF Early PostDoc Mobility fellowship (P2EZP3_148773) and a FWF Erwin Schrödinger fellowship (J3774-B25). CS was funded by Convocatoria para proyectos de investigación en Ciencias Básicas (COLCINECIAS)-2014, contract No. FP44842-103-2015. WJP was supported by an MRC Centenary Award.

